# Turnip mosaic virus-based gRNA delivery system for plant genome editing

**DOI:** 10.64898/2026.04.22.720221

**Authors:** Ekkachai Khwanbua, Ryan R. Lappe, Austin A. Bierl, Steven A. Whitham

## Abstract

Plant virus-based gRNA delivery systems offer a rapid alternative to stable transformation for CRISPR-mediated genome editing, but potyvirus-based platforms in Cas9-expressing plants are still underexplored. Here, we developed a turnip mosaic virus (TuMV)-based system for gRNA delivery in Cas9-expressing *Nicotiana benthamiana* and tested whether Csy4-mediated gRNA processing could improve editing efficiency. A TuMV construct carrying a gRNA targeting *PHYTOENE DESATURASE* (*NbPDS*) induced detectable editing in both infiltrated and systemic tissues, although editing frequencies were low. Incorporation of the bacterial endoribonuclease Csy4 increased editing efficiencies in the two *NbPDS* genes, raising editing in infiltrated leaves to 7.1–13.8% for *NbPDSa* and 7.6–23.0% for *NbPDSb*, while lower but reproducible editing was detectable in systemic leaves. The TuMV-Csy4 platform also supported editing of a second endogenous target, *MAGNESIUM CHELATASE SUBUNIT H* (*NbChlH*), and enabled multiplex editing of *NbPDS* and *NbChlH* regardless of guide order. Editing efficiencies were consistently higher in infiltrated leaves than in systemic leaves, and no visible photobleaching or chlorosis was observed in systemic tissues despite confirmed molecular editing. To assess the potential for heritable editing, a tRNA^Ile^ mobility element was fused to the *NbPDS* gRNA. Although this construct increased somatic editing, no albino progeny were recovered after screening approximately 20,000 seedlings, indicating that heritable editing was not achieved under these conditions. Together, these results establish TuMV as a platform for Cas9-based gRNA delivery and show that Csy4-mediated processing improves editing efficiency, supports multiplex targeting, and demonstrates the feasibility of potyvirus-based genome editing systems in plants.

## 1 INTRODUCTION

Genetic modification can be achieved through both random and targeted (site-specific) mutagenesis. Although several methods have been used to induce random mutations, such as chemical mutagenesis, physical mutagenesis by radiation, and biological mutagenesis by transposon or T-DNA in plants, isolating the desired mutations from random ones is extremely difficult (Alonso & Ecker, 2006). The advent of genome editing technologies has enabled precise gene editing, offering new possibilities for crop improvement and gene function analysis (Zhang *et al*., 2021). The fundamental approach to genome editing involves using a sequence-specific nuclease to create a DNA double-strand break (DSB) at a specific target site. This break can then be repaired through either the non-homologous end-joining pathway (NHEJ) or the donor-dependent homology-directed repair (HDR) pathway, resulting in genetic modifications (Doudna & Charpentier, 2014). Early sequence-specific nucleases, such as meganucleases (Puchta *et al*., 1993), zinc-finger nucleases (Wright *et al*., 2005), and transcription activator-like effector nucleases (Christian *et al*., 2010), were devised to achieve target-specific mutagenesis based on the principles of DNA-protein recognition. However, difficulties in protein design, synthesis, and validation continue to hinder the widespread adoption of these engineered nucleases for routine use.

CRISPR (Clustered Regulatory Interspaced Short Palindromic Repeats)/Cas (CRISPR associated) is an RNA-mediated adaptive phage immunity system in archaea and bacteria (Mali *et al*., 2013). This system utilizes DNA–RNA recognition and binding to achieve sequence-specific cleavage of nucleic acids, allowing for the simple programming of CRISPR/Cas9 and other CRISPR/Cas systems to introduce DSBs at desired target sites. The *Streptococcus pyogenes* Type II CRISPR/Cas9 (*Sp*Cas9) system was the first shown to precisely cleave DNA both *in vitro* and *in vivo* (Jinek *et al*., 2012; Gasiunas *et al*., 2012; Mali *et al*., 2013). The CRISPR/Cas9 system for gene editing comprises two key components: the Cas9 nuclease and a single guide RNA (gRNA), which is an artificial fusion of a crRNA and a transactivating crRNA (tracrRNA). The gRNA and Cas9 protein form a Cas9/gRNA complex, with the gRNA specifying the target DNA site. This complex facilitates the cleavage of DNA sequences adjacent to 5’-NGG-3’ protospacer-adjacent motifs (PAMs) (Chen *et al*., 2019). However, one of the major drawbacks of CRISPR/Cas9-mediated genome editing is its dependence on plants that can be transformed and regenerated, a process that is labor-intensive and time-consuming (Mao *et al*., 2019). To broaden the application of genome editing systems to plants recalcitrant to transformation or to accelerate the process in existing systems, new technologies are needed to overcome the challenges associated with delivering genome editing reagents *in planta*.

An alternative approach to transformation and regeneration for introducing genetic elements into plants is the use of plant viruses as delivery platforms. Numerous plant viruses have been engineered for biotechnological applications, including somatic and heritable genome editing through virus-induced gene editing (VIGE) (Oh *et al*., 2021). Several plant RNA virus-based vectors, such as tobacco rattle virus (TRV) (Ali *et al*., 2015), tobacco mosaic virus (TMV) (Cody *et al*., 2017), pea early-browning virus (PEBV) (Ali *et al*., 2018), beet necrotic yellow vein virus (BNYVV) (Jiang *et al*., 2019), foxtail mosaic virus (FoMV) (Mei *et al*., 2019a), potato virus X (PVX) (Uranga *et al*., 2021), barley stripe mosaic virus (BSMV) (Tamilselvan-Nattar-Amutha *et al*., 2023), tomato spotted wilt virus (TSWV) (Liu *et al*., 2023), and tobacco ringspot virus (TRSV) (Yoshida *et al*., 2024) have been developed to deliver gRNAs systemically in Cas9-expressing plants, facilitating targeted genome editing in model or crop plants. While some virus-based vectors used for VIGE exhibit relatively broad host ranges, many remain restricted to particular taxonomic groups, and platforms capable of reliable systemic delivery across a broad range of crop species are still limited (Oh *et al*., 2021).

The *Potyviridae* family is the largest group of plant-infecting RNA viruses and collectively infects over 30 plant families, including many economically important crop species (Walsh & Jenner, 2002; Wylie *et al*., 2017). Several recombinant potyviruses have been developed for gene overexpression (Kelloniemi *et al*., 2006; Chen *et al*., 2007; Majer *et al*., 2017; Mei *et al*., 2019b; Achs *et al*., 2022) and gene silencing (Xie *et al*., 2021; Tuo *et al*., 2021; Wang *et al*., 2025). Recent work has shown that potyvirus-based vectors can enable virus-induced genome editing in Cas12a-expressing plants (Merwaiss *et al*., 2026). However, the application of potyvirus-based systems in Cas9-expressing plants and strategies to improve gRNA processing and editing efficiency are still underexplored. Here, we investigated the potential of a TuMV-based platform for delivering functional gRNAs in Cas9-expressing *Nicotiana benthamiana* and demonstrate that incorporation of the RNA endoribonuclease Csy4 enhances editing efficiency, enables multiplex targeting, and facilitates the investigation of heritable genome editing.

## 2 MATERIALS AND METHODS

### 2.1 Plant growth conditions and viral inoculation

Transgenic *N. benthamiana* plants expressing Cas9 derived from *S. pyogenes* (*Sp*Cas9) were germinated on LC1 Grower’s Mix (Sungro) in a greenhouse at 22-25°C. The plants were subjected to a 16-h light (320–350 μmol m^2^/s) and 8-h dark photoperiod with 47%-55% relative humidity and were fertilized weekly with Peter’s Excel 15-15-15 (ICL Performance Products). Four-week-old *N. benthamiana* plants were infiltrated with *Agrobacterium tumefaciens* (strain GV3101) harboring viral vectors suspended in an infiltration buffer consisting of 10 mM MgCl_2_, 10 mM MES, and 200 mM acetosyringone (pH 5.6) (Mei *et al*., 2019a).

### 2.2 Constructions of TuMV-derived vectors

The turnip mosaic virus (TuMV) genome is a positive-sense single-stranded RNA of approximately 10 kilobases (kb) that encodes ten proteins from a single open reading frame (ORF), as well as the PIPO protein, which is translated from a short ORF embedded within the P3 cistron of the polyprotein (Chung *et al*., 2008). Conventional cloning sites in potyvirus vectors are typically positioned between cistrons, taking advantage of natural and engineered protease cleavage sites within the single ORF for heterologous expression. However, cloning sites within the viral ORF are unsuitable for insertion of the tracrRNA sequence, because such an insertion would introduce stop codons and possibly frameshifts into the viral genome.

To generate a TuMV vector capable of delivering gRNA for genome editing, the gRNA cassette was inserted immediately after the stop codon following the coat protein (CP) cistron and upstream of the 3′ untranslated region (3′ UTR) (Figure 1A). All TuMV constructs generated in this study were derived from pCB-TuMV-GFP (UK1 strain; GenBank: EF028235.1). The GFP coding region in pCB-TuMV-GFP was excised and replaced with a multiple cloning site (MCS) designated MCS2. An additional *Pvu*II restriction site was introduced immediately after the polyprotein stop codon to create TuMV-MCS2-3′UTR. TuMV-MCS2-3′UTR was then digested with *Pvu*II, and the tracrRNA sequence was inserted to produce TuMV-MCS2-3′UTRt (Data S1). Finally, a crRNA sequence targeting *N. benthamiana PHYTOENE DESATURASE* (*NbPDS*; Niben101Scf01283g02002.1 and Niben101Scf14708g00023.1) (Beernink *et al*., 2022) was cloned into TuMV-MCS2-3′ UTRt to generate TuMV:gNbPDS.

**Figure 1.**
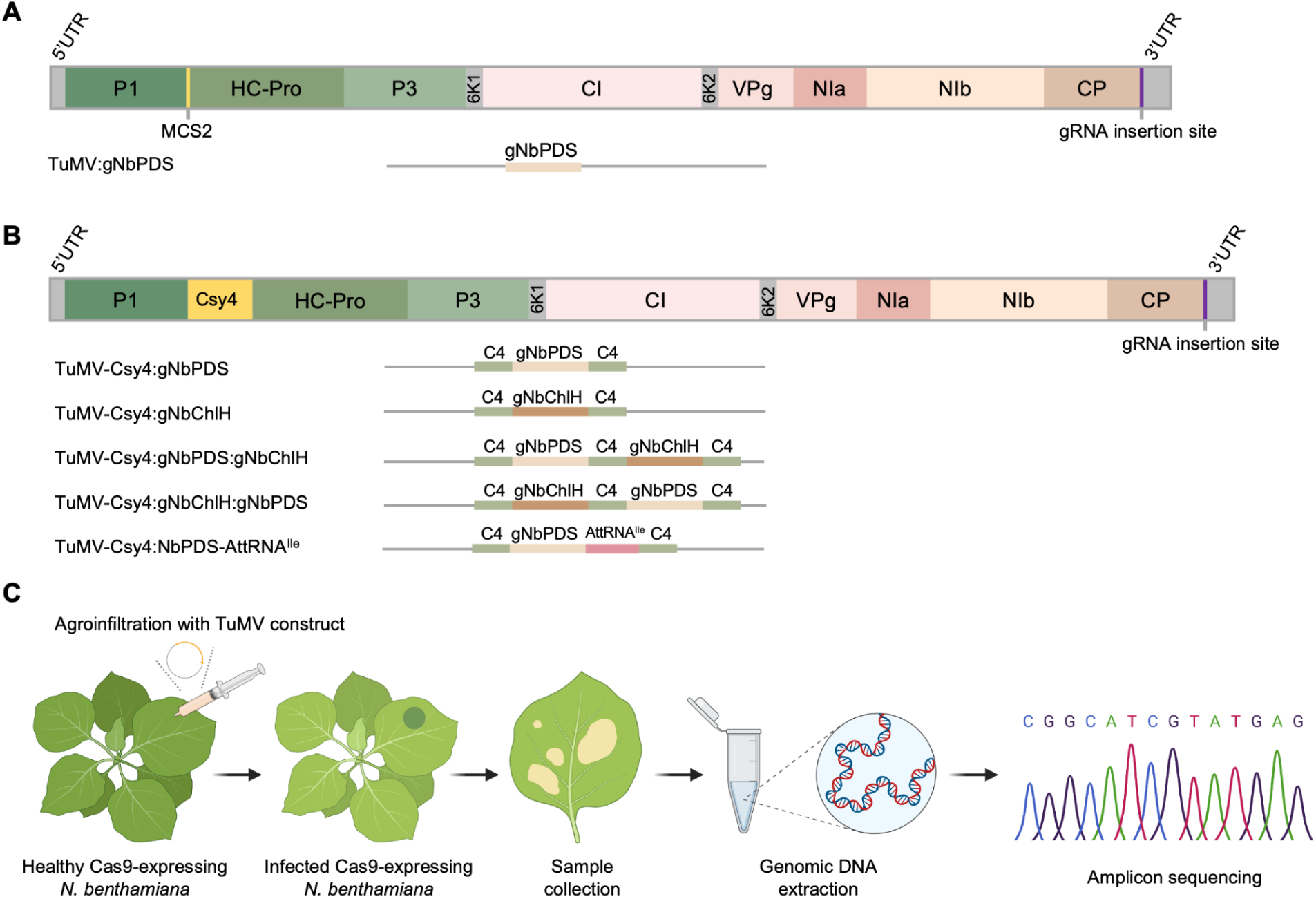
Schematic overview of TuMV-mediated genome editing in *N. benthamiana*. (A) Schematic representation of the TuMV genome showing the positions of major cistrons (P1, HC-Pro, P3, 6K1, CI, 6K2, VPg, NIa, NIb, and CP) and the insertion site used for gRNA expression. (B) TuMV-Csy4-based vector architecture. The Csy4 processing sequence was inserted between P1 and HC-Pro to enable precise cleavage and release of individual gRNAs. Representative constructs carrying gRNAs targeting *N. benthamiana PHYTOENE DESATURASE* (*NbPDS*) and/or *N. benthamiana MAGNESIUM CHELATASE SUBUNIT H* (*NbChlH*) are shown as examples. C4 indicates the position of the Csy4 cleavage sites. (C) Overall workflow of TuMV-mediated genome editing. Cas9-expressing *N. benthamiana plants* (*NbCas9*) were agroinfiltrated with TuMV-based vectors carrying gRNA expression cassettes. Following systemic infection, genomic DNA was extracted from infiltrated and systemic leaves for PCR amplification and mutation analysis by restriction enzyme digestion and sequencing. Figure 1C was created in BioRender (https://BioRender.com/pj3jxlk).

To develop a Csy4-processing TuMV-based gRNA delivery system, TuMV-MCS2-3′UTR was digested with *Pvu*II, and synthetic double-stranded DNA oligonucleotides were ligated to introduce an *Sfo*I restriction site, generating TuMV-MCS2-3′SfoI. The Csy4 coding sequence was amplified from PGK1p-Csy4-pA (Construct 2; Addgene plasmid #55196; RRID:Addgene_55196) (Nissim *et al*., 2014) using primers Csy4HiFi-F and Csy4HiFi-R (Table S1), and the resulting amplicon was cloned into TuMV-MCS2-3′SfoI to yield TuMV-Csy4-3′SfoI. TuMV-Csy4-3′SfoI was subsequently digested with *Sfo*I and assembled with the tracrRNA sequence containing Csy4 recognition (C4) sites to generate TuMV-Csy4-3′t. crRNA sequences targeting *N. benthamiana PHYTOENE DESATURASE* (*NbPDS*) and *MAGNESIUM CHELATASE SUBUNIT H* (*NbChlH*; Niben101Scf04388g00011.1), or both genes, were then cloned into TuMV-Csy4-3′t to construct TuMV-Csy4:gNbPDS, TuMV-Csy4:gNbChlH, TuMV- Csy4:gNbPDS:gNbChlH, TuMV-Csy4:gNbChlH:gNbPDS, and TuMV-Csy4:gNbPDS-AttRNA^Ile^ (Figure 1B).

### 2.3 Reverse transcription PCR (RT-PCR) analysis

RT-PCR was performed to assess the systemic viral infection and the stability of gRNAs. Leaf samples were collected from *N. benthamiana* plants at 35 days post inoculation (dpi). Total RNA was isolated from approximately 50 mg of leaf tissue using TRIzol Reagent (Thermo Fisher Scientific, Waltham, MA, USA), and 2 µg of RNA was reverse transcribed using the Maxima First Strand cDNA Synthesis Kit (Thermo Fisher Scientific, Waltham, MA, USA). TuMV was detected by RT-PCR using primers TuMV9606F and TuMV9879R (Table S1). Primers NbPP2A-F and NbPP2A-R (Table S1) were used as internal reference controls for *N. benthamiana* (Ellison *et al*., 2020).

### 2.4 Verification of mutations in *N. benthamiana* genomic DNA

gRNAs for *NbPDS* and *NbChlH* were designed to target sequences containing restriction enzyme recognition sites located 3-4 bp upstream of PAMs, enabling the detection of gene edits through restriction enzyme digestion of PCR amplicons, as previously described (Mei *et al*., 2019a). Inoculated leaves were collected at 7 dpi, and systemic leaves were sampled at 35 dpi. Genomic DNA was extracted from leaf tissue using the cetyltrimethylammonium bromide (CTAB) method (Murray & Thompson, 1980). Amplicons spanning the gRNA target sequence were generated using Platinum SuperFi II PCR Master Mix (Thermo Fisher Scientific, Waltham, MA, USA) with primers NbPDSa-F and NbPDSa-R for *NbPDSa*, NbPDSb-F and NbPDSb-R for *NbPDSb*, and NbChlH-F and NbChlH-R for *NbChlH* (Table S1). *NbPDS* and *NbChlH* amplicons were digested with *Nco*I and *Mlu*CI and resolved on 1% or 2% agarose gels, respectively. In a separate analysis, PCR amplicons were purified and subjected to Sanger sequencing, and gene editing efficiency was estimated using the Tracking of Indels by DEcomposition (TIDE) algorithm by comparing sequence chromatograms of edited samples to those of non-edited control plants (Brinkman *et al*., 2014).

Heritable mutations were assessed by collecting seeds from parent plants and screening them for the presence of albino seedlings after germination. Biallelic mutations in *NbPDS* result in an albino phenotype in *N. benthamiana* seedlings, which was visually evaluated on LC1 Grower’s Mix (Sungro). The number of seedlings screened was estimated based on seed weight (1,000 seeds per 78.4 mg) (Beernink *et al*., 2022).

## 3 RESULTS

### 3.1 TuMV-mediated editing of *NbPDS* in *Nb*Cas9 plants

The overall workflow of TuMV-mediated gene editing is shown in Figure 1C, illustrating agroinfiltration of *N. benthamiana* Cas9 plants (*Nb*Cas9) with TuMV-based vectors, followed by genomic DNA extraction and mutation analysis. Using this approach, we used a well-established gRNA targeting the *N. benthamiana PHYTOENE DESATURASE* (*NbPDS*) gene with an overlapping *Nco*I recognition site to create TuMV:gNbPDS. Genomic DNA was extracted from infiltrated leaves at 7 days post-inoculation (dpi) and from systemic leaves at 35 dpi to determine if mutations occurred in *NbPDSa* or *NbPDSb*. Polymerase chain reaction (PCR) amplified 858 bp and 845 bp fragments spanning the target sites in *NbPDSa* and *NbPDSb*, respectively, which were subsequently digested with *Nco*I. PCR products from non-infected *Nb*Cas9 (Figure 2A,S1A) and TuMV:EV-infected *Nb*Cas9 (Figure 2B,S1B) negative control plants yielded two digestion products for each target, 373 and 485 bp for *NbPDSa*, and 372 and 473 bp for *NbPDSb* (Figure 2I). In contrast, TuMV:gNbPDS-infected *Nb*Cas9 plants (Figure 2C,S1C) yielded additional undigested fragments of 858 bp for *NbPDSa* and 845 bp for *NbPDSb* at both 7 (Figure S2A) and 35 dpi (Figure 2I), indicating mutations in the targeted sequences. Editing efficiencies in infiltrated leaves ranged from 0.8-1.3% for *NbPDSa* (Figure 3A) and 2.1-2.7% for *NbPDSb* (Figure 3B), whereas lower editing efficiencies of 0.8-1.2% for *NbPDSa* (Figure 3C) and 0.9-1.4% for *NbPDSb* (Figure 3D) were observed in systemic leaves.

**Figure 2.**
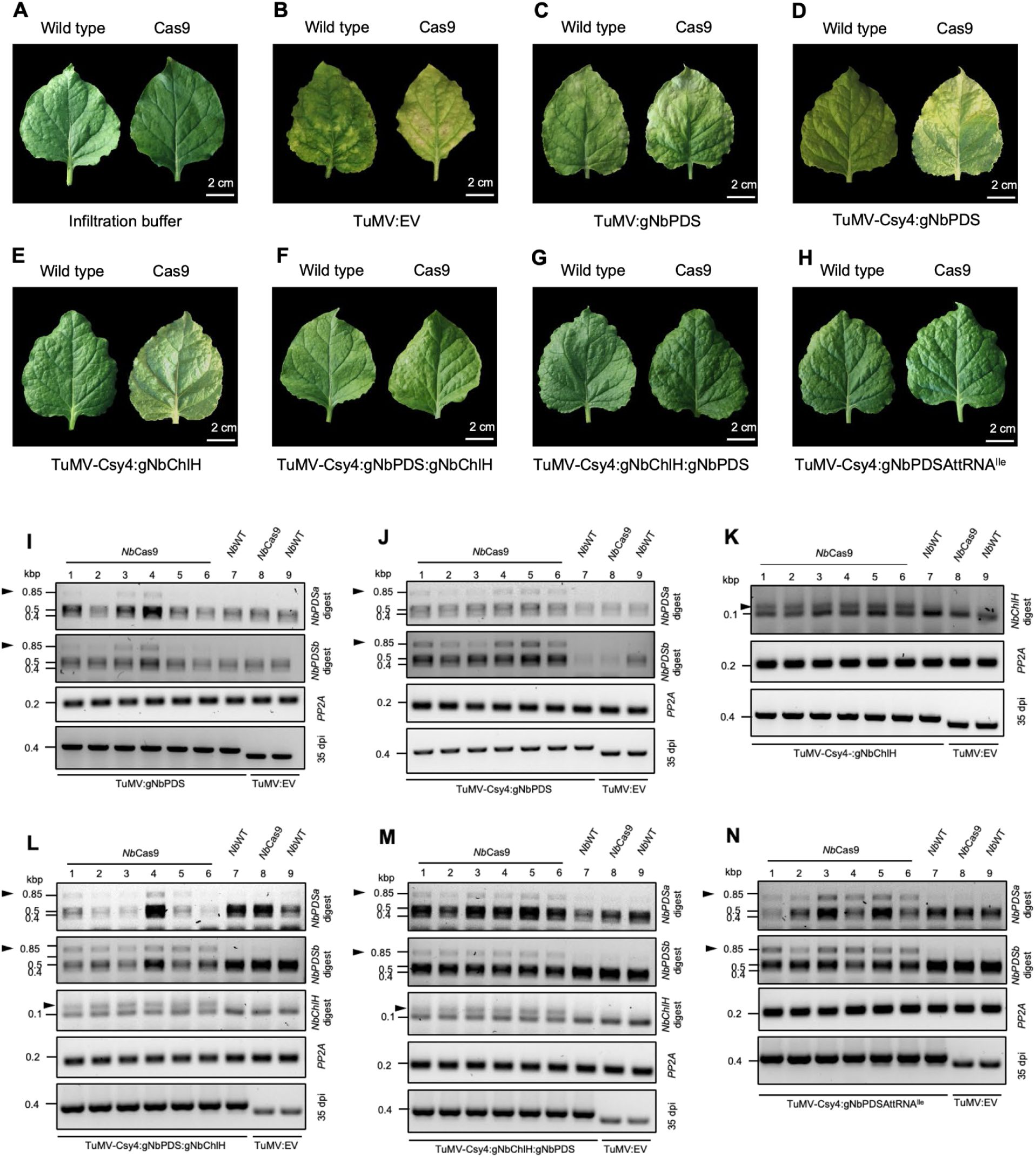
TuMV-mediated genome editing in Cas9-expressing *N. benthamiana*. (A–H) Representative systemic leaves collected at 35 days post-inoculation (dpi) from wild-type and *NbCas9* plants agroinfiltrated with infiltration buffer (A), TuMV:EV (B), TuMV:gNbPDS (C), TuMV-Csy4:gNbPDS (D), TuMV-Csy4:gNbChlH (E), TuMV- Csy4:gNbPDS:gNbChlH (F), TuMV-Csy4:gNbChlH:gNbPDS (G), and TuMV-Csy4:gNbPDSAttRNA^Ile^ (H). Scale bars = 2 cm. (I–N) Restriction digestion analysis of genomic DNA amplicons spanning the target sites in systemic leaves collected at 35 dpi from *Nb*Cas9 plants infected with TuMV:gNbPDS (I), TuMV-Csy4:gNbPDS (J), TuMV-Csy4:gNbChlH (K), TuMV-Csy4:gNbPDS:gNbChlH (L), TuMV-Csy4:gNbChlH:gNbPDS (M), and TuMV-Csy4:gNbPDSAttRNA^Ile^ (N). *NbPDSa* and *NbPDSb* amplicons were digested with *Nco*I, and *NbChlH* amplicons were digested with *Mlu*CI. Undigested bands indicate targeted editing events. *NbPP2A* was amplified as a *N. benthamiana* control. The presence of each viral construct at 35 dpi was assessed by RT-PCR, with TuMV:EV included as a control for comparison with gRNA-containing constructs.

**Figure 3.**
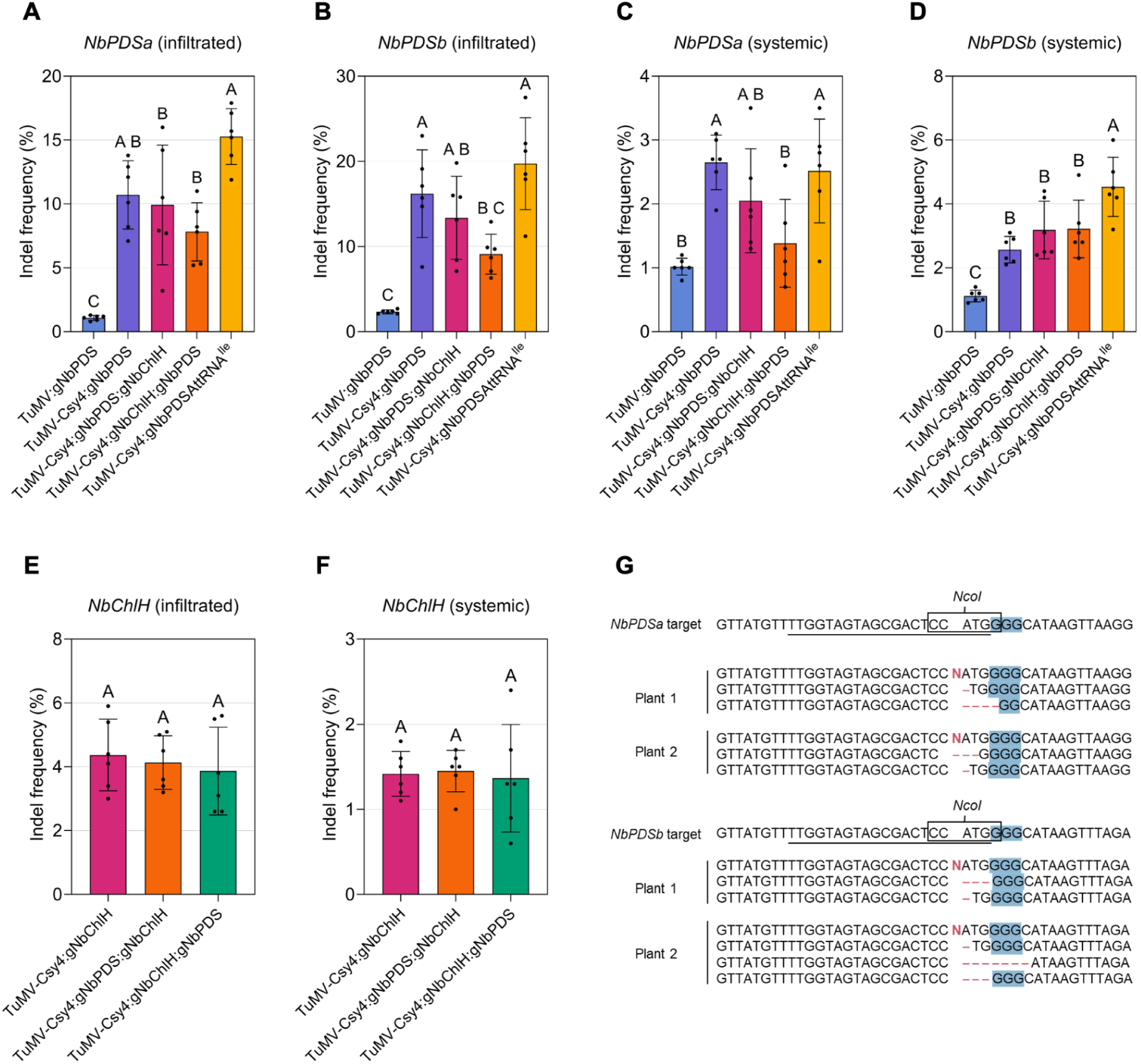
Editing efficiencies of TuMV-based genome editing constructs in Cas9-expressing *N. benthamiana*. (A–D) Indel frequencies at the *NbPDSa* and *NbPDSb* loci in infiltrated leaves collected at 7 days post-inoculation (dpi) and systemic leaves collected at 35 dpi from *Nb*Cas9 plants infected with TuMV:gNbPDS, TuMV-Csy4:gNbPDS, TuMV-Csy4:gNbPDS:gNbChlH, TuMV-Csy4:gNbChlH:gNbPDS, and TuMV-Csy4:gNbPDSAttRNA^Ile^. Indel frequencies were determined by TIDE analysis of Sanger sequencing chromatograms. (A) *NbPDSa* in infiltrated leaves. (B) *NbPDSb* in infiltrated leaves. (C) *NbPDSa* in systemic leaves. (D) *NbPDSb* in systemic leaves. (E–F) Indel frequencies at the *NbChlH* locus in infiltrated and systemic leaves from *Nb*Cas9 plants infected with TuMV-Csy4:gNbChlH, TuMV-Csy4:gNbPDS:gNbChlH, and TuMV-Csy4:gNbChlH:gNbPDS. (E) *NbChlH* in infiltrated leaves. (F) *NbChlH* in systemic leaves. (G) Representative indel sequences detected at the *NbPDSa* and *NbPDSb* target sites in systemic leaves of *Nb*Cas9 plants infected with TuMV-Csy4:gNbPDSAttRNA^Ile^. The gRNA target sequence is underlined, PAM sequences are highlighted in blue, the *Nco*I recognition site is boxed, and insertions or deletions are indicated in red. Bars represent mean ± SE. Statistical significance was assessed by ordinary one-way ANOVA followed by Tukey’s multiple comparisons test. Different letters indicate significant differences among treatments at *P* < 0.05; treatments sharing the same letter are not significantly different.

### 3.2 Csy4-mediated gRNA processing improves TuMV-based genome editing efficiency

Although TuMV:gNbPDS induced targeted gene editing in *Nb*Cas9 at a low frequency and caused noticeable infection symptoms, it did not induce the photobleaching phenotype associated with PDS loss-of-function mutations in *Nb*Cas9 (Qin *et al*., 2007). This observation prompted us to test whether optimizing gRNA processing with Csy4 could enhance editing efficiency. Csy4 is an endoribonuclease from a bacterial CRISPR-Cas system that cleaves precursor CRISPR RNA transcripts into individual CRISPR RNAs, and it has been adapted in plants to facilitate the release of mature gRNAs for Cas9, thereby improving editing efficiency (Luo *et al*., 2024). Accordingly, TuMV-Csy4:gNbPDS was developed to test this strategy. Editing efficiency was assessed using the same PCR-*Nco*I digest assay described above. Undigested amplicons from TuMV-Csy4:gNbPDS-infected *Nb*Cas9 plants (Figure 2D,S1D), corresponding to 858 bp (*NbPDSa*) and 845 bp (*NbPDSb*), were detected at both 7 (Figure S2B) and 35 dpi (Figure 2J) and appeared more abundant than in TuMV:gNbPDS-infected samples (Figure 2I,S2A). Editing efficiencies in infiltrated leaves increased to 7.1–13.8% for *NbPDSa* (Figure 3A) and 7.6–23% for *NbPDSb* (Figure 3B), whereas lower efficiencies of 1.9–3.1% for *NbPDSa* (Figure 3C) and 2.1–3.1% for *NbPDSb* (Figure 3C) were observed in systemic leaves. These plants exhibited undigested fragments in both infiltrated and systemic leaves, indicating that Csy4-mediated processing enhanced gRNA maturation and editing efficiency during systemic viral spread. Despite this increase in editing efficiency, no visible photobleaching phenotype was observed in systemic leaves at 35 dpi (Figure 2D).

### 3.3 Csy4-mediated TuMV system supports editing of multiple endogenous loci

To determine if TuMV-Csy4 can target additional endogenous loci in *N. benthamiana*, we designed a gRNA targeting the *MAGNESIUM CHELATASE SUBUNIT H* (*NbChlH*) gene. *NbChlH* encodes a subunit of magnesium chelatase required for chlorophyll biosynthesis, and its disruption produces a characteristic chlorotic phenotype, making it a common marker for genome editing in *N. benthamiana* (Senthil-Kumar & Mysore, 2014). The selected gRNA overlapped an *Mlu*CI recognition site, allowing restriction digestion-based detection of editing events. We constructed TuMV-Csy4:gNbChlH and tested whether it could induce mutations at this second endogenous target. Similar to *NbPDS*, editing at the *NbChlH* locus was assessed by PCR amplification of a 115-bp fragment spanning the target site followed by *Mlu*CI digestion. TuMV-Csy4:gNbChlH-infected *Nb*Cas9 plants (Figure 2E, S1E) contained undigested fragments in both infiltrated (Figure S2C) and systemic leaves (Figure 2K), indicating that Csy4-mediated processing enabled efficient targeting of the *NbChlH* locus across infected tissues. Editing efficiencies in infiltrated leaves ranged from 3.0-5.9% (Figure 3E), whereas systemic leaves showed lower efficiencies of 1.1-1.8% (Figure 3F). Despite confirmed editing at both stages, no visible chlorotic or white patch phenotype was observed in systemic leaves at 35 dpi (Figure 2E).

### 3.4 Multiplex genome editing using the TuMV-Csy4 system

Having demonstrated that Csy4 expression from TuMV enhances editing of *NbPDS* and *NbChlH* individually, we next examined whether the system could also support multiplex genome editing. Multiplex editing, in which different gRNAs are expressed simultaneously, has been shown to broaden the scope and enhance the efficiency of genetic engineering (Lowder *et al*., 2015). To explore this possibility, we tested the co-delivery of two gRNAs in *N. benthamiana* using TuMV. Specifically, we multiplexed the gRNAs targeting *NbPDS* and *NbChlH*, generating TuMV-Csy4:gNbPDS:gNbChlH and TuMV-Csy4:gNbChlH:gNbPDS. Editing efficiency was evaluated at 7 dpi and 35 dpi using the same PCR-*Nco*I and PCR-*Mlu*CI digest assays described above. Undigested fragments corresponding to *NbPDSa* (858 bp), *NbPDSb* (845 bp), and *NbChlH* (115 bp) were detected for both constructs at both time points (Figure 2L–M; S2D–E), confirming editing at all three target sites. As expected, editing efficiencies were consistently higher in infiltrated leaves than in systemic leaves for both constructs, with *NbPDS* targets exhibiting greater editing frequencies than *NbChlH*. In infiltrated tissues, *NbPDSa* and *NbPDSb* editing reached up to 20% (Figure 3A,B), whereas *NbChlH* editing was lower (∼3-6%) (Figure 3E). In systemic leaves, editing efficiencies were uniformly reduced across all targets, ranging from ∼0.6% to ∼4.9% (Figure 3C,D,F). Consistent with the low editing frequencies, no photobleaching or chlorosis was observed in systemic leaves of plants infected by the multiplex constructs at 35 dpi (Figure 2F–G; S1F–G).

### 3.5 tRNA^Ile^ fusion enhances somatic editing but not does not enable heritable edits

We were also interested in whether adding a tRNA^Ile^ sequence could increase the frequency of editing in systemic leaves and the likelihood of heritable editing by promoting gRNA mobility. Certain tRNA molecules, including tRNA^Ile^, act as mobile RNA signals that enable fused gRNAs to move into meristematic cells, where heritable mutations can be initiated (Ellison *et al*., 2020). To test this concept in a TuMV-based system, we engineered TuMV-Csy4:NbPDSAttRNA^Ile^, in which an *Arabidopsis thaliana* tRNA^Ile^ element was fused to the 3’ end of the NbPDS gRNA. PCR amplicons spanning the target site were digested with *Nco*I, and undigested fragments corresponding to *NbPDSa* (858 bp) and *NbPDSb* (845 bp) were detected at both 7 dpi (Figure S2F) and 35 dpi (Figure 2N), confirming successful somatic editing at both stages. Editing efficiencies reached 11.9-17.9% for *NbPDSa* (Figure 3A,G) and 11.2-27.5% for *NbPDSb* (Figure 3B,G) in infiltrated leaves, whereas systemic leaves showed lower efficiencies of 1.1-3.5% (Figure C) and 3.2-6.0% (Figure 3D), respectively. No visible photobleaching or other phenotypes were detected in systemic leaves at 35 dpi (Figure 2H,S1H). To determine whether edits were transmitted to the next generation, approximately 20,000 progeny were screened for albino phenotypes as an indicator of germline editing. TuMV-Csy4:NbPDSAttRNA^Ile^-infected *Nb*Cas9 plants did not produce any albino progeny, indicating that the addition of tRNA^Ile^ was insufficient to promote heritable editing under these conditions.

## 4 DISCUSSION

In light of these findings, our work can be positioned within recent advances in potyvirus-based VIGE systems. Potyviruses have been engineered for genome editing, although existing platforms differ in nuclease class, vector design, and biological outcomes. A recent study investigated gRNA delivery in *Lb*Cas12a-expressing plants using tobacco etch virus (TEV) as the primary vector and subsequently extended to TuMV and lettuce mosaic virus (LMV) (Merwaiss *et al*., 2026). TEV enabled efficient editing but caused severe symptoms in *N. benthamiana*, requiring attenuation of the silencing suppression activity of HC-Pro to enable plant survival and seed production, whereas TuMV achieved editing in most samples and outperformed LMV. In our system, severe symptoms were observed only in plants infected with TuMV:gNbPDS, whereas Csy4-containing constructs did not show similarly severe symptoms and were able to produce seeds (Figure S1). Our study extends potyvirus-based genome editing into a *Sp*Cas9 context and introduces an alternative strategy for gRNA processing within a TuMV backbone.

A key distinction between the two systems lies in gRNA processing. In the Cas12a-based platform, intrinsic RNase activity enables crRNA processing and simplifies guide design, whereas Cas9-based systems may benefit more from additional guide RNA processing strategies to generate functional gRNAs. Consistent with this difference, incorporation of Csy4 in our TuMV platform was associated with increased editing at the *NbPDS* locus relative to the non-Csy4 construct, particularly in infiltrated tissue, while editing was also detectable in systemic leaves (Figure 3A–D). These results suggest that efficient gRNA processing may be an important factor influencing Cas9-based editing in viral systems. Previous studies have also shown that Csy4-mediated processing of guide RNAs can improve Cas9 editing in plant virus-based platforms, including apple latent spherical virus (ALSV)- and TRV-based systems (Luo *et al*., 2021, 2024).

Beyond single-locus editing, the TuMV-Csy4 platform supported targeting of a second endogenous locus, *NbChlH*, and enabled multiplex editing of *NbPDS* and *NbChlH* in both guide orders (Figure 3). Editing was consistently higher in infiltrated leaves than in systemic leaves, indicating reduced editing efficiency in systemic tissues in this platform. Although these efficiencies were sufficient for molecular detection, they were not high enough to produce visible photobleaching or chlorosis in systemic leaves. These observations indicate that maintaining editing efficiency during systemic infection remains a challenge.

The heritability results illustrate both the potential and the current limitation of this platform. Fusion of AttRNA^Ile^ increased somatic editing of *NbPDS*, but no albino progeny were recovered among ∼20,000 seedlings, suggesting that this mobility element was not sufficient to promote detectable germline editing under our conditions. These findings suggest that, although the system can mediate somatic editing, efficient editing in meristematic or germline cells may still be limited. Previous VIGE studies have shown that heritable editing can be improved by mobile guide modules in some systems, but it is still influenced by the viral vector, guide design, host background, and access to meristematic tissues (Oh *et al*., 2021; Nagalakshmi *et al*., 2022; Weiss *et al*., 2025).

Overall, this study defines a distinct contribution to the expanding potyviral VIGE toolbox. Our results show that TuMV can be engineered for Cas9-based gRNA delivery, that Csy4 improves editing efficiency in this context, and that the vector can accommodate both multiplex guide arrays and RNA mobility elements. At the same time, the relatively low systemic editing efficiencies and absence of heritable editing indicate that further optimization is needed, particularly in gRNA processing, accumulation, and access to meristematic tissues. Future work should focus on alternative guide-processing strategies, insertion sites within the TuMV genome, mobility elements with stronger germline activity, and regeneration-based recovery of edited lines. Taken together, these results support TuMV as a promising foundation for the development of more versatile potyvirus-based editing platforms in Cas9-expressing plants.

## AUTHOR CONTRIBUTIONS

EK, RRL, and SAW conceived and designed the study. EK conducted the experiments and drafted the manuscript. RRL and AAB assisted with the experiments. SAW edited the manuscript with input from all coauthors. The final version was approved by all authors.

## ACKNOWLEDGMENTS

We thank the ISU DNA Facility for providing instrumentation and technical expertise. This work was supported by the ISU Plant Sciences Institute, the ISU Crop Bioengineering Center, the Iowa Soybean Association, and USDA NIFA Hatch Project IOW04308. EK was supported by an Anandamahidol fellowship from the government of Thailand.

## DATA AVAILABILITY STATEMENT

The data generated in this study are included in the article and its supporting information. Additional data are available from the corresponding author upon request.

## SUPPORTING INFORMATION

**Figure S1.**
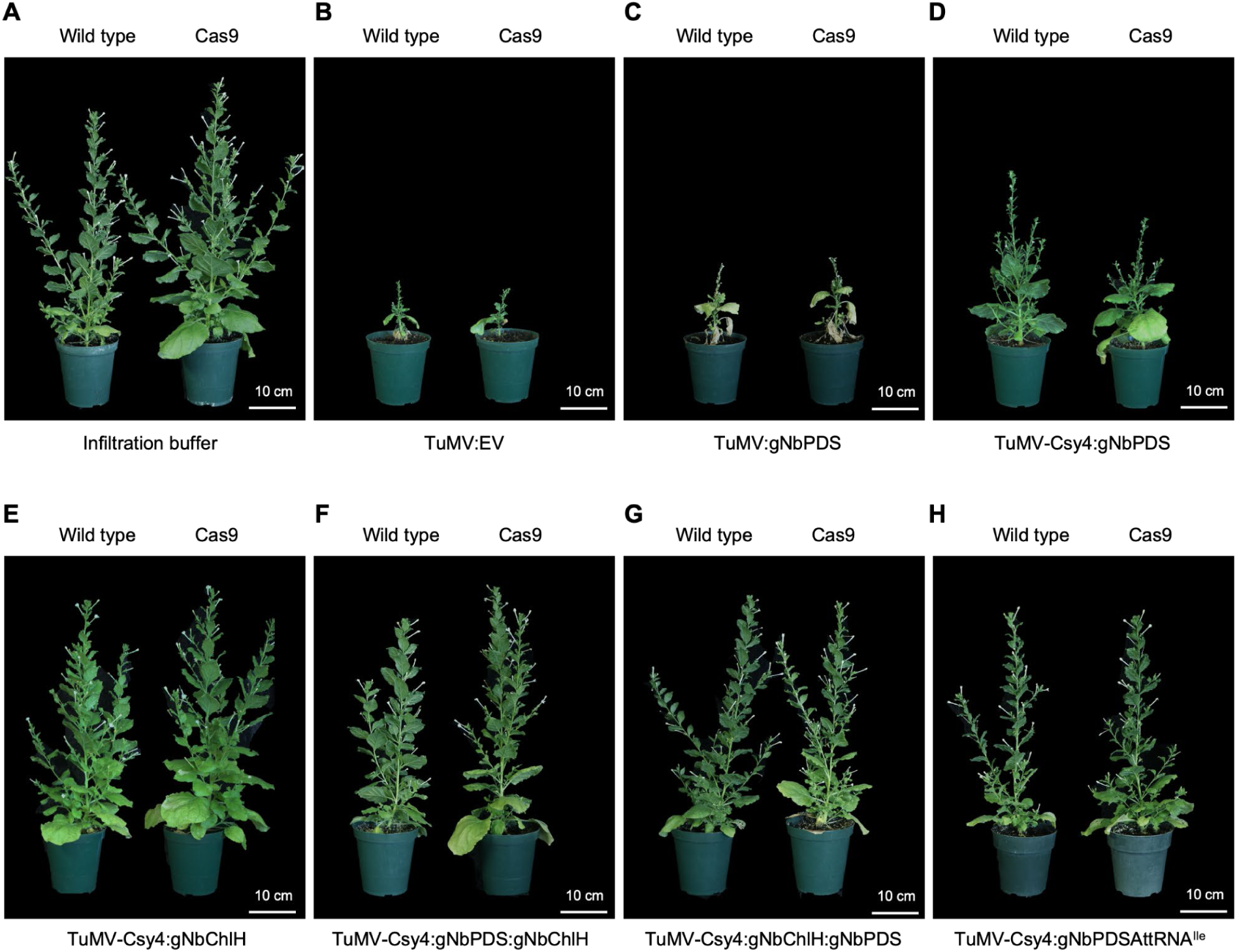
Whole-plant phenotypes of TuMV-infected *N. benthamiana* and Cas9-expressing *N. benthamiana* (*NbCas9*) plants. (A–H) Wild-type and *NbCas9* plants were infected with buffer, TuMV:EV, TuMV:gNbPDS, TuMV-Csy4:gNbPDS, TuMV-Csy4:gNbChlH, TuMV-Csy4:gNbPDS:gNbChlH, TuMV-Csy4:gNbChlH:gNbPDS, or TuMV-Csy4:gNbPDSAttRNA^Ile^ and photographed at 35 days post-inoculation (dpi). Scale bars = 10 cm.

**Figure S2.**
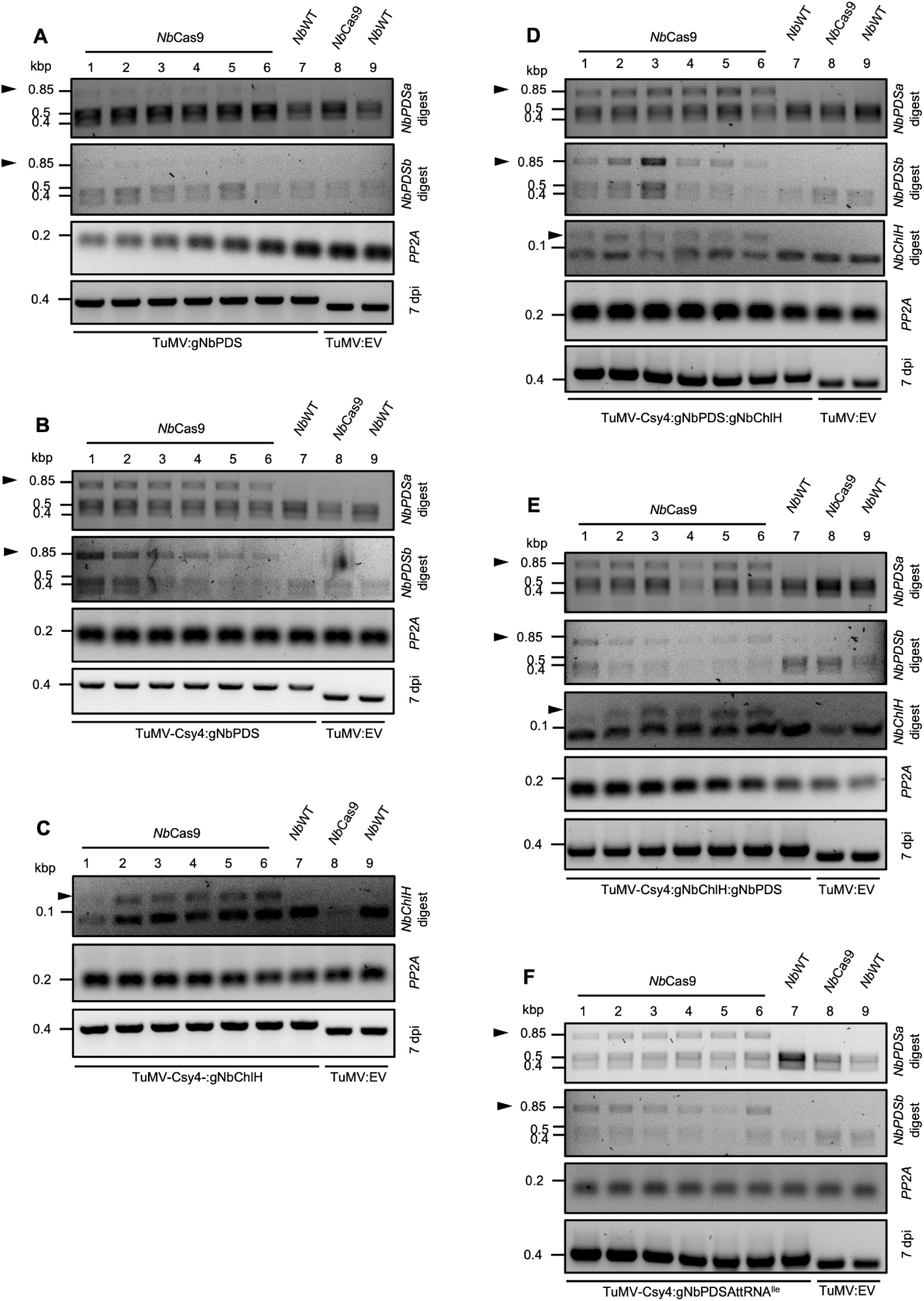
PCR-restriction assays showing genome editing in infiltrated *N. benthamiana* leaves at 7 days post-inoculation (dpi). (A–F) DNA was extracted from infiltrated leaves and analyzed for editing at the *NbPDS* (*Nco*I) and *NbChlH* (*Mlu*CI) loci. Amplicons were digested with *Nco*I and/or *Mlu*CI, depending on the targeted construct, to detect loss of restriction sites caused by sequence changes at the target regions. For *NbPDS*, 858 bp (*NbPDSa*) and 845 bp (*NbPDSb*) fragments were digested with *Nco*I. In unedited samples, digestion produced 373 + 485 bp (*NbPDSa*) and 372 + 473 bp (*NbPDSb*) fragments, whereas undigested 858 bp or 845 bp bands (arrow) indicated successful editing. For *NbChlH*, a 115 bp fragment containing the *Mlu*CI site was analyzed. In unedited samples, digestion produced 84 bp and 31 bp fragments, while undigested 115 bp products (arrows) confirmed editing at the *ChlH* locus.

**Table S1.**
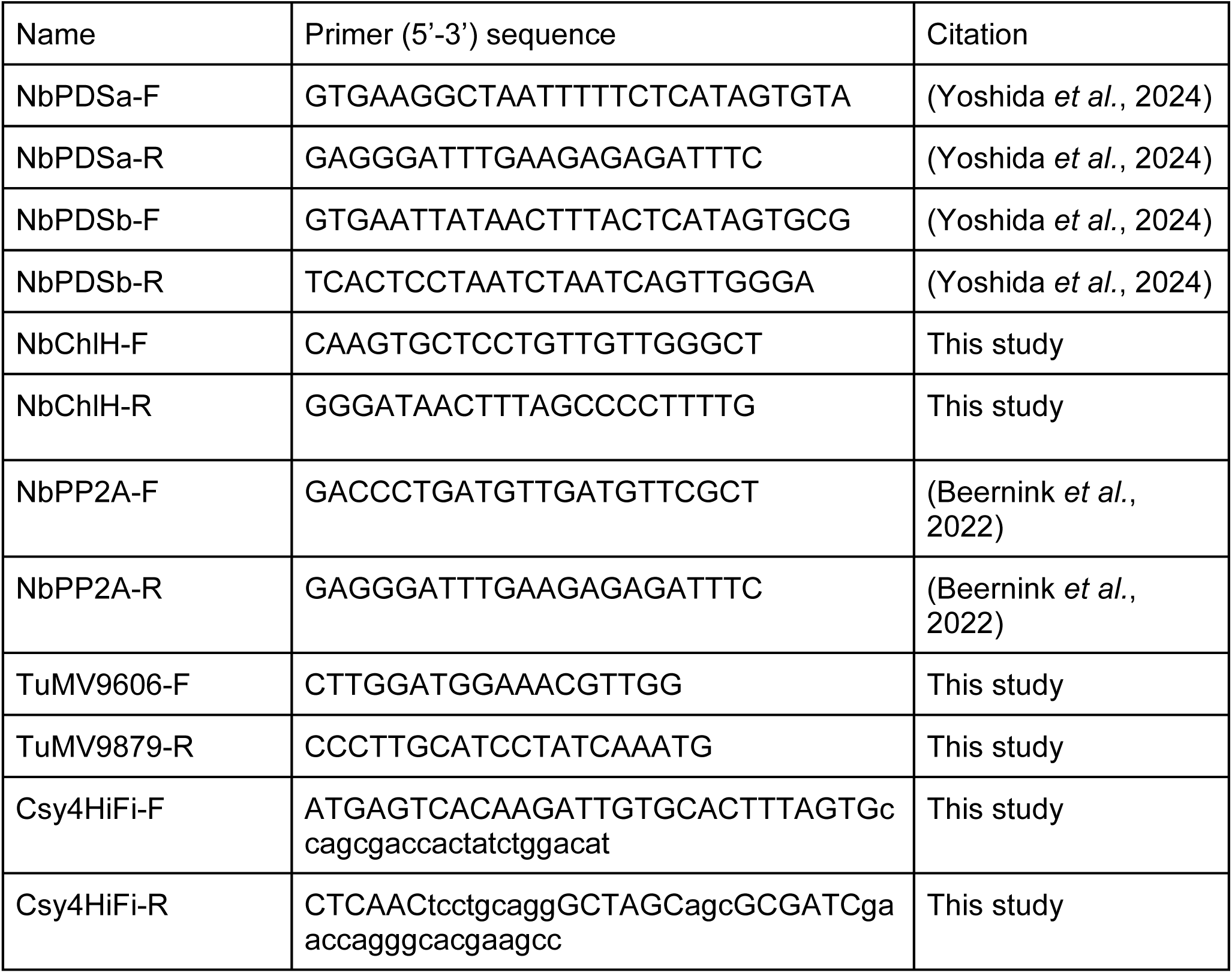
List of oligonucleotide primers used in this study with sequences.

**Data S1. Genome sequence of TuMV-3UTRt.**

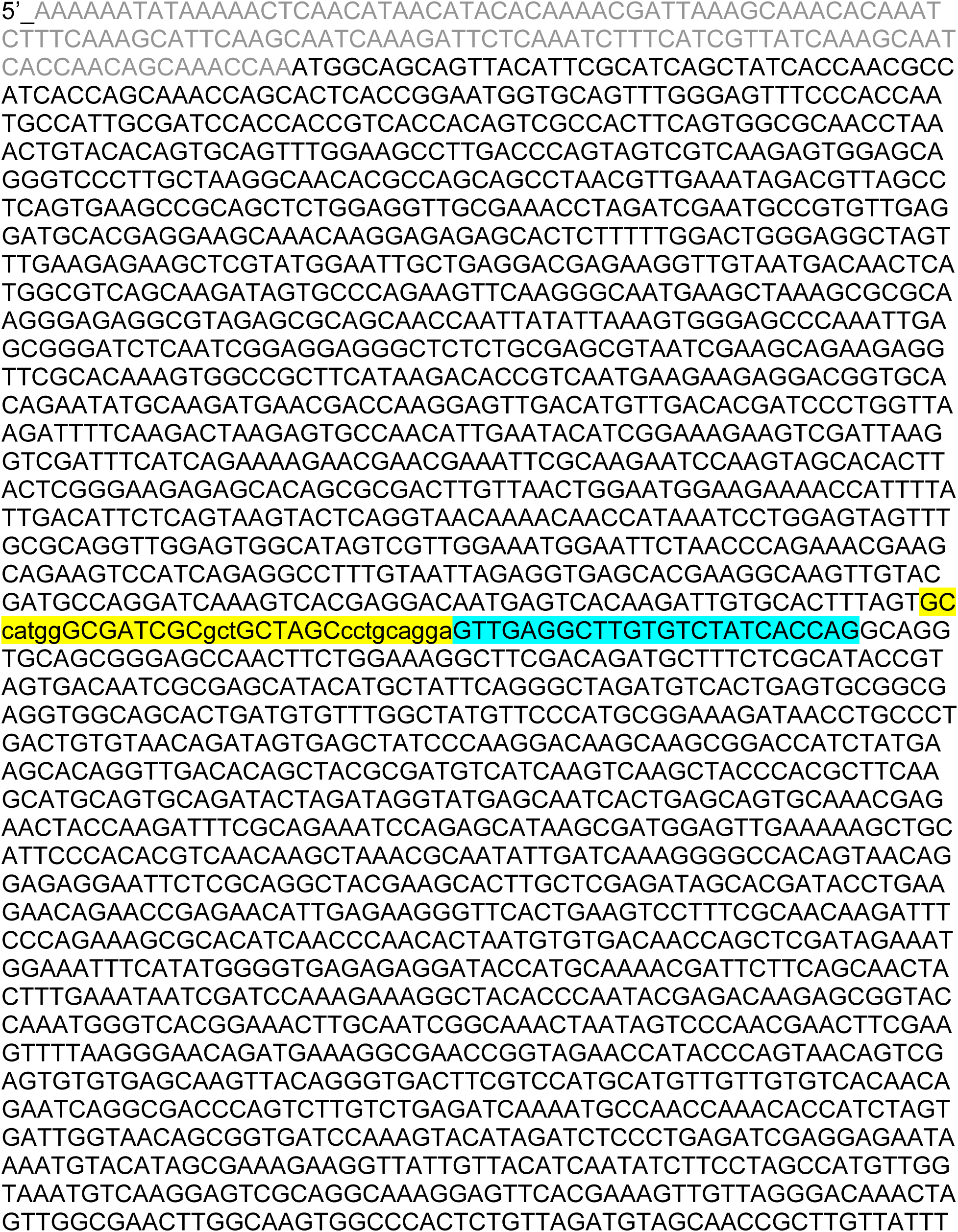
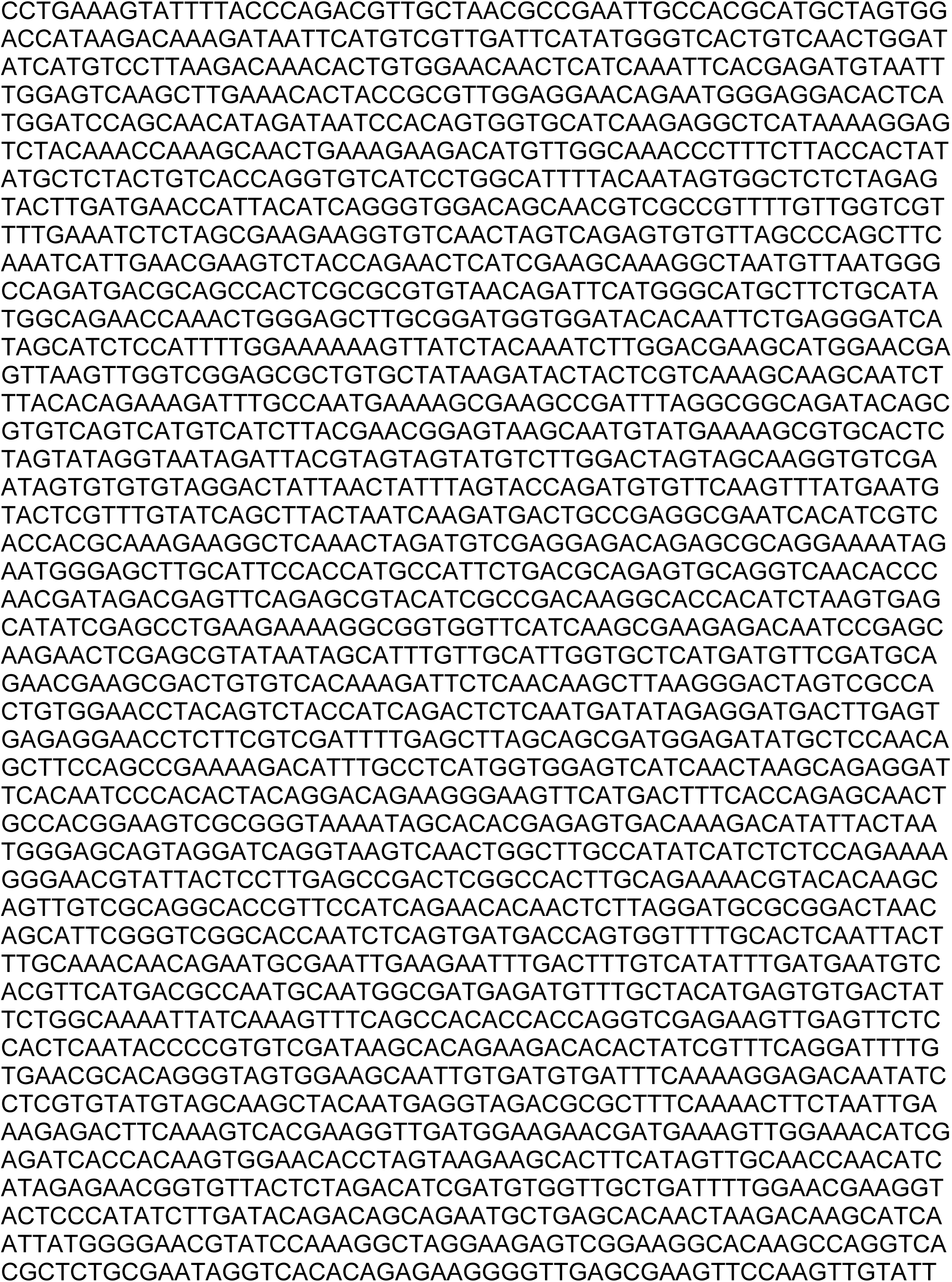
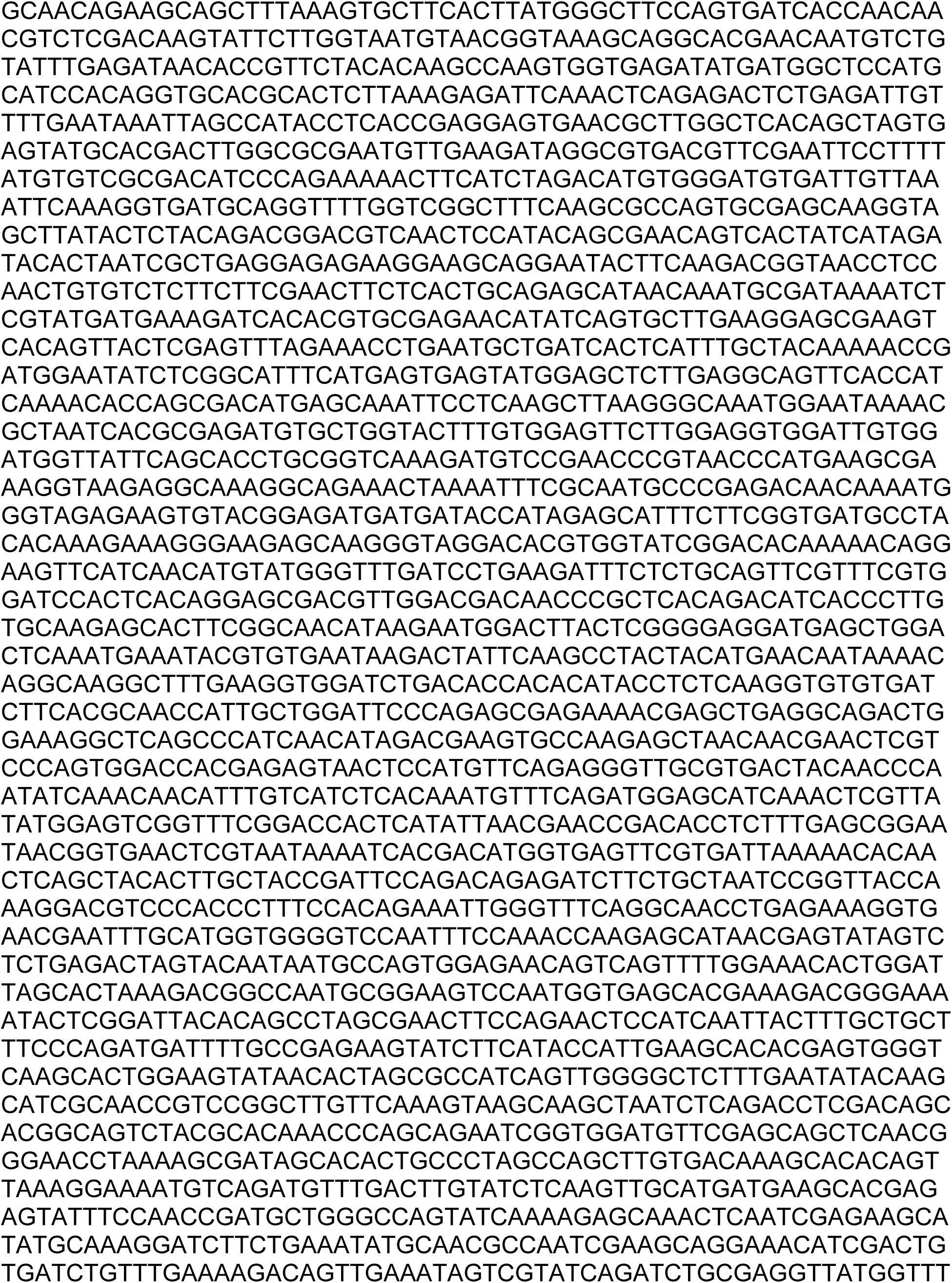
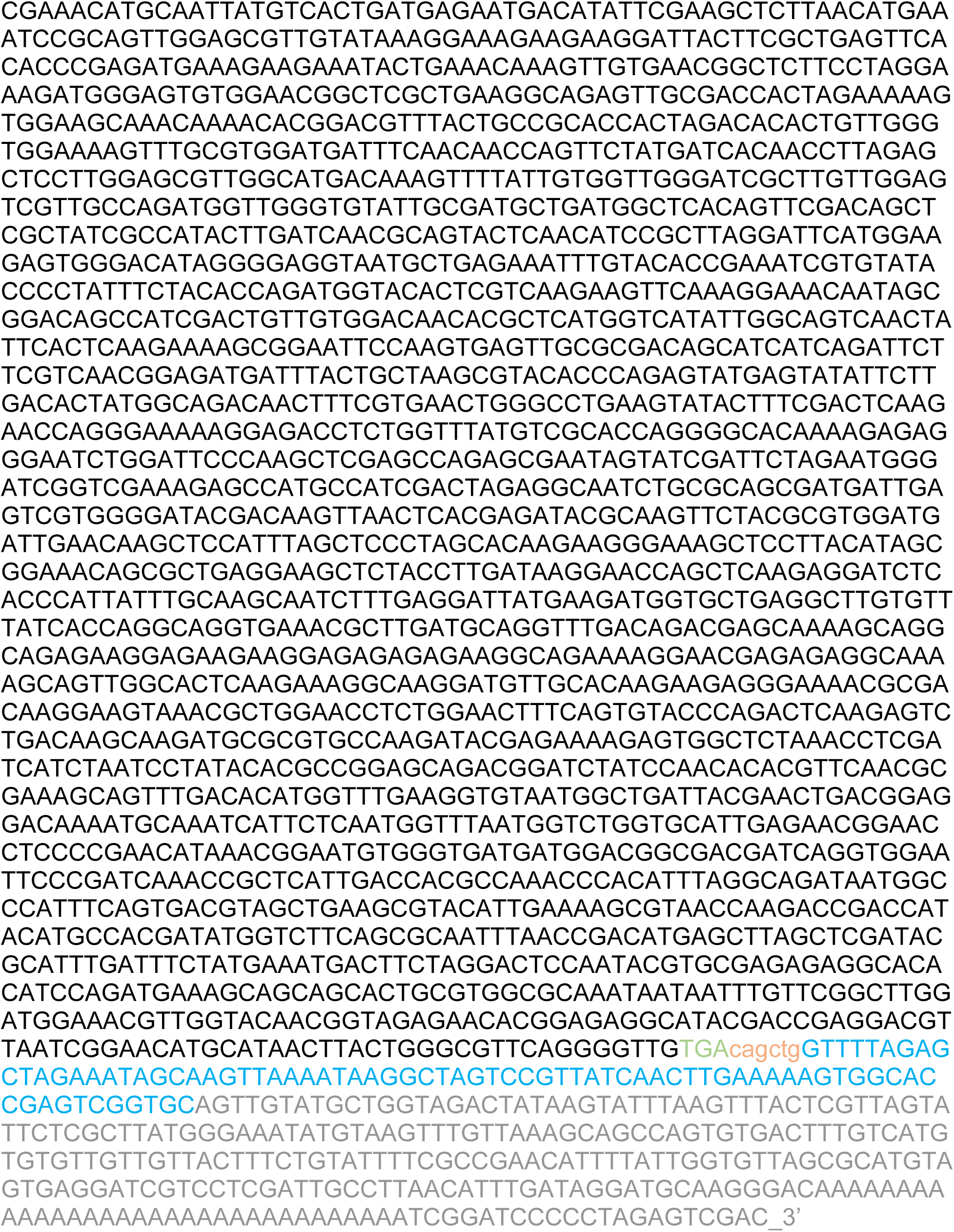
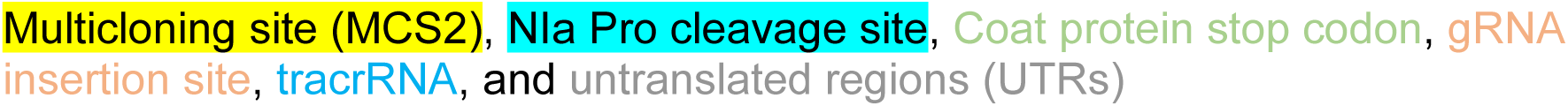

